# Discovery of Bavachin as a Dual Trk-A/Trk-B Agonist via Integrated Computational Screening and Experimental Validation

**DOI:** 10.64898/2026.09.10.750759

**Authors:** Sachin P. Patil, Bella Kuehn, Ryan Schlosser, John Samohod

## Abstract

Neurotrophins regulate neuronal survival, differentiation, and synaptic plasticity through activation of tropomyosin receptor kinase (Trk) receptors, and their dysregulation is strongly implicated in neurodegenerative disorders, including Alzheimer’s disease. However, the therapeutic application of recombinant neurotrophins remains limited by poor bioavailability, rapid degradation, and inadequate blood–brain barrier permeability. Here, we report the discovery of Bavachin as a dual Trk-A/Trk-B neurotrophin mimetic via an integrated computational screening and experimental validation approach. Structure-based virtual screening of a bioactive natural product library derived from Traditional Chinese Medicine, combined with a novel consensus-ranking strategy, enabled the identification of flavonoid scaffolds targeting neurotrophin-binding sites of Trk receptors. Among these, Bavachin demonstrated robust, dose-dependent activation of both Trk-A and Trk-B receptors, but not Trk-C, indicating receptor selectivity. Induced-fit docking and molecular dynamics simulations revealed stable binding of Bavachin within conserved and receptor-specific ligand-binding pockets, consistent with its dual agonist activity. Collectively, this study identifies Bavachin as a putative small-molecule neurotrophin mimetic and establishes an integrated computational–experimental framework for the discovery of dual Trk-A/Trk-B agonists for neurodegenerative disease therapeutics.

## 1. Introduction

Neurotrophins are a family of growth factors that play a central role in neuronal survival, differentiation, synaptic plasticity, and repair of the nervous system [1–3]. The major neurotrophins, nerve growth factor (NGF), brain-derived neurotrophic factor (BDNF), neurotrophin-3 (NT-3), and neurotrophin-4/5 (NT-4/5), exert their biological effects primarily through activation of the tropomyosin receptor kinase (Trk) family of receptor tyrosine kinases, namely Trk-A, Trk-B, and Trk-C [1,4]. Ligand binding to the extracellular domain of Trk receptors induces receptor dimerization and autophosphorylation, triggering downstream signaling cascades such as PI3K/Akt, MAPK/ERK, and PLCγ pathways, which collectively regulate neuronal growth and survival [2,5].

Dysregulation of neurotrophin–Trk signaling has been strongly implicated in a wide range of neurological and neurodegenerative disorders, including Alzheimer’s disease, Parkinson’s disease, depression, and peripheral neuropathies [6–8]. In particular, reduced BDNF/Trk-B signaling has been associated with synaptic dysfunction and cognitive decline in Alzheimer’s disease, while impaired NGF/Trk-A signaling contributes to degeneration of cholinergic neurons [6,9]. These findings have established Trk receptors as attractive therapeutic targets for neurodegenerative diseases.

Despite the clear therapeutic potential of neurotrophins, their clinical application has been limited by several pharmacological challenges, including poor blood–brain barrier (BBB) permeability, short half-life, and pleiotropic effects [10,11]. To overcome these limitations, significant efforts have been directed toward the development of small-molecule neurotrophin mimetics capable of selectively activating Trk receptors [12–14]. Such compounds offer advantages in terms of oral bioavailability, metabolic stability, and central nervous system (CNS) penetration. Despite representing a promising alternative, however, identification of such small-molecule neurotrophin mimetics has proven challenging. While numerous Trk inhibitors have been successfully developed, discovery of small-molecule agonists capable of activating Trk receptors remains limited.

Among small-molecule candidates, flavonoids and other natural products have attracted considerable attention due to their structural diversity and neuroprotective properties [15–17]. One of the most widely studied compounds, 7,8-dihydroxyflavone (7,8-DHF), has been reported to act as a Trk-B agonist and to exhibit neuroprotective effects in multiple disease models [13,18,19]. However, recent studies have raised questions regarding its direct mechanism of Trk-B activation, suggesting that its effects may be indirect or context-dependent [20,21]. This controversy highlights a broader challenge in the field– the identification of true small-molecule agonists that can reproducibly and directly activate Trk receptors.

Structurally, activation of Trk receptors is mediated by complex interactions between neurotrophin ligands and the extracellular immunoglobulin-like domains of the receptors [22–24]. Crystallographic studies of Trk–neurotrophin complexes have revealed conserved and receptor-specific binding interfaces that govern ligand recognition and receptor activation [22,23]. Based on these structural insights, we hypothesized that small molecules capable of engaging key residues within these binding pockets may mimic neurotrophin-induced receptor activation. However, rational identification of such molecules remains challenging, as conventional structure-based virtual screening approaches often preferentially identify inhibitors rather than agonists [25].

Nevertheless, natural product databases, particularly those derived from traditional medicine systems, represent a rich source of bioactive scaffolds for drug discovery [26,27]. Notably, compounds present in bioavailable fractions of traditional herbal formulations are more likely to possess favorable pharmacokinetic properties, making them attractive candidates for CNS drug development [27]. Integrating such databases with advanced computational screening strategies has the potential to enable the discovery of novel small-molecule Trk agonists.

In this context, our present study employed an integrated computational and experimental approach to identify small-molecule neurotrophin mimetics targeting Trk-A and Trk-B receptors. By leveraging structural insights from Trk–neurotrophin complexes, we developed a novel consensus-ranking strategy to prioritize compounds capable of engaging conserved and receptor-specific binding pockets on Trk-A and Trk-B receptors. Screening of a bioactive natural product database derived from the Traditional Chinese Medicine (TCM) led to the identification of flavonoid-based compounds as promising candidates. Experimental validation using cell-based reporter assays demonstrated that Bavachin induces robust, dose-dependent activation of both Trk-A and Trk-B receptors, but not Trk-C. Complementary induced-fit docking and molecular dynamics simulations further supported stable binding of Bavachin within key receptor pockets, consistent with its dual agonist activity. Collectively, these findings identify Bavachin as a potential small-molecule neurotrophin mimetic and provide a generalizable framework for the discovery of dual Trk-A/Trk-B agonists for neurodegenerative disease therapeutics.

## 2. Results

The present study involved Integrated virtual and experimental screening of a database containing drug-like compounds derived from the TCM, viz. Database of Constituents Absorbed into the Blood and Metabolites of TCM (DCABM-TCM) [28]. The natural compounds have been the basis of 3/4^th^ of approved drugs worldwide during the past half century, and as such, are a precious source of novel chemical scaffolds for modern drug discovery [26]. Specifically, the DCABM-TCM database may prove to be a great resource of active natural products for the central nervous system (CNS) drug development [29,30], offering potential dual Trk-A/B agonists that are both bioactive and bioavailable. Therefore, virtual screening hits from the DCABM-TCM database were subjected to both experimental Trk activation studies and further computational analyses including the induced-fit docking (IFD) and molecular dynamics (MD) simulation studies.

### 2.1. Structural basis of neurotrophin recognition by Trk-A and Trk-B receptors

To identify potential small-molecule agonists capable of activating Trk receptors, we first analyzed the published X-ray crystal structures of the extracellular domains of Trk-A bound to nerve growth factor (NGF) (PDB ID: 1WWW) [22] and Trk-B bound to neurotrophin-4/5 (NT-4/5) (PDB ID: 1HCF) [23]. These structures show that receptor activation occurs through ligand-induced dimerization in which two neurotrophin molecules bridge two receptor ectodomains, generating a symmetric complex resembling a “bat”, in which the neurotrophin dimer forms the central body and the two Trk receptor molecules extend outward as wings (**Figure 1**).

**Figure 1.**
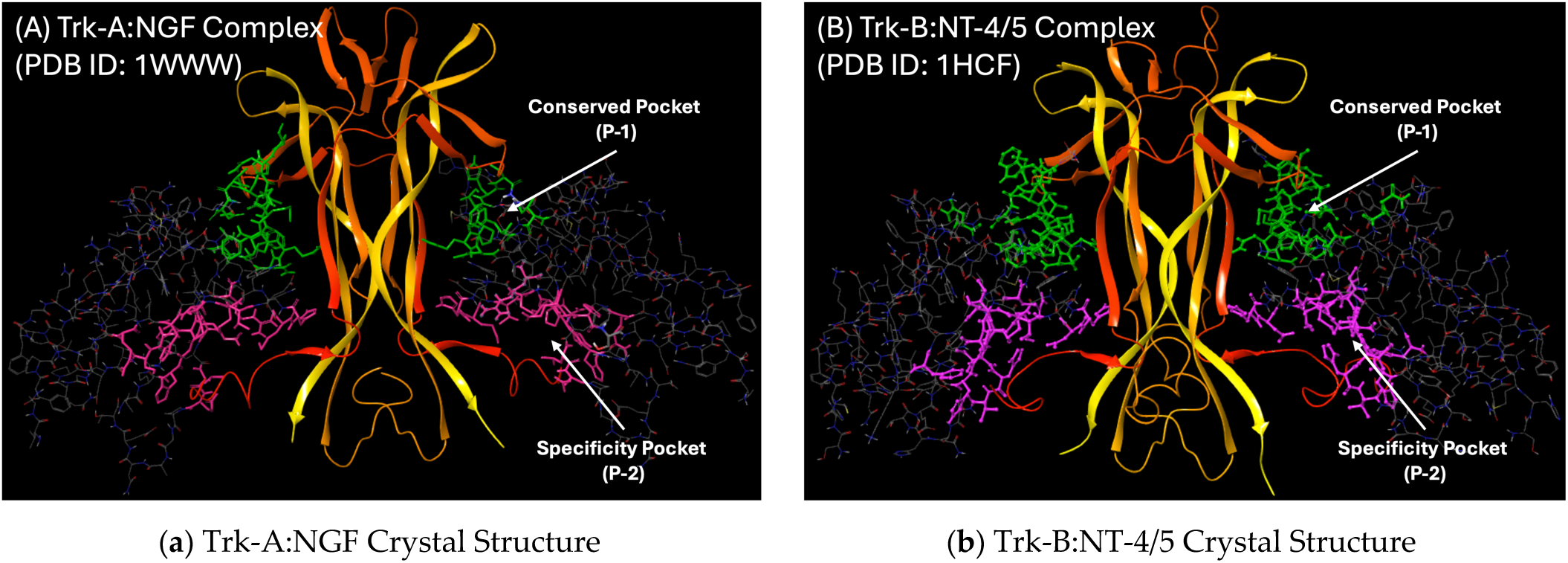
The crystal Structures of (A) Trk-A and (B) Trk-B receptors (stick models) in complex with their endogenous neurotrophin ligands, NGF and NT-4/5 (ribbon models), respectively.

Inspection of both structures revealed the presence of two distinct ligand-binding pockets on each receptor. The first site, referred to here as the conserved pocket (Pocket-1; P-1), corresponds to a structurally conserved interface present in both Trk-A and Trk-B that mediates interactions between the central β-sheet region of the neurotrophin ligand and residues from the immunoglobulin-like domains of the Trk receptors. Structural studies have shown that residues such as Asn-349, Gln-350, and Phe-327 in Trk-A, together with interacting residues from NGF including His-84 and Arg-103, contribute to this conserved binding interface [22]. Similarly, in the Trk-B:NT-4/5 complex, analogous residues including Asp-349 and Asn-350 mediate key hydrogen-bonding and π-π stacking interactions that stabilize ligand binding [23]. The conservation of these residues and their structural arrangement across Trk receptors explains the cross-reactivity observed among neurotrophins and suggests the possibility of targeting this pocket with small molecules capable of activating multiple Trk receptors.

In contrast, the second region, termed the specificity pocket (Pocket-2; P-2), displays notable structural and sequence divergence between Trk family members. In Trk-A, this pocket forms a hydrophobic canyon defined in part by residues such as Val294, Leu333, Ser304, and Arg347, with contributions from surrounding structural elements including a disulfide-linked region that shapes the pocket architecture. In the Trk-B complex, different residues define this region, such as His299, Asp298, and His343, underlying the receptor-specific interactions observed with individual neurotrophins. These observations highlight the potential of exploiting both conserved and specificity pockets to identify selective or dual Trk agonists.

### 2.2. Druggability analysis of Trk-A and Trk-B neurotrophin-binding pockets

To determine whether the identified pockets are suitable for small-molecule targeting, we performed druggability analysis using the DoGSiteScorer server [31]. Both conserved and specificity pockets on Trk-A and Trk-B were predicted to be druggable binding sites (**Figure 2**). The Trk-A specificity pocket displayed the highest druggability score (0.60), whereas the conserved pockets of both receptors showed moderate but significant scores, indicating favorable cavity geometry and physicochemical characteristics for ligand binding. Importantly, these predicted druggable pockets coincided with the experimentally observed binding sites of NGF and NT-4/5 in the crystal structures, suggesting that small molecules targeting these sites may mimic neurotrophin-induced receptor activation. Here, Pocket-1 on both Trk-A and Trk-B was predicted to contain two sub-pockets in both receptors, providing additional structural features potentially suitable for accommodating diverse chemical scaffolds. These analyses thus supported the suitability of these pockets for structure-based virtual screening to identify candidate Trk agonists from natural product libraries.

**Figure 2.**
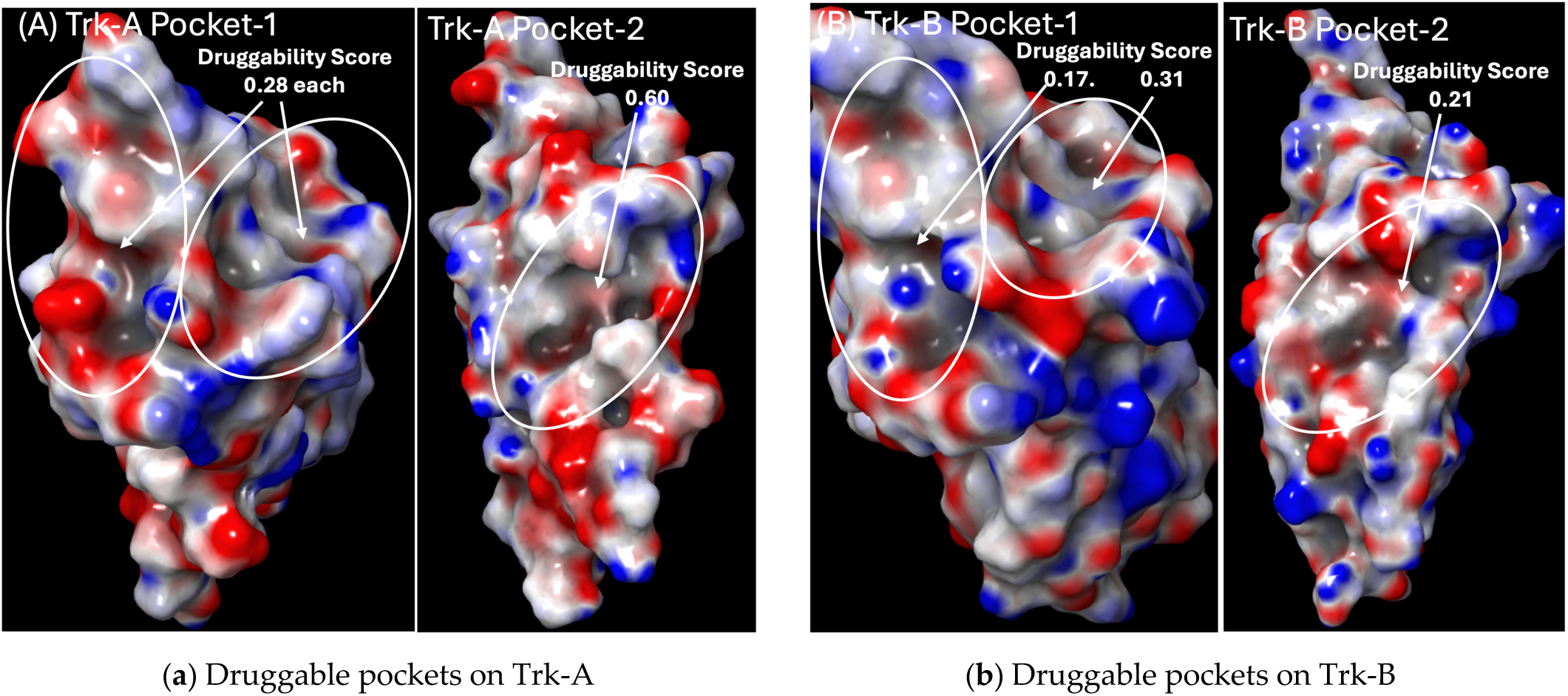
The druggable binding sites on (A) Trk-A and (B) Trk-B receptors.

### 2.3. Structure-based virtual screening and development of a consensus-ranking strategy

Using the identified pockets, a virtual screening campaign was carried out against the DCABM-TCM database of bioactive natural compounds. To validate the docking protocol, previously reported Trk antagonists (from crystal structures with high resolution of ≤ 2Å) as well as published agonists were included in the screening dataset. Docking was performed against four receptor sites: Trk-A-P1, Trk-A-P2, Trk-B-P1, and Trk-B-P2. The AutoDock Vina [32] docking algorithm was used, wherein the receptor remains rigid while the ligands are flexibly docked. Interestingly, several known Trk inhibitors ranked highly in individual pocket dockings (**Table 1**), with multiple compounds appearing within the top 100 hits for at least one pocket (≤ 100 ranks shown in the bold).

**Table 1.** AutoDock Vina individual and consensus docking ranks of known Trk antagonists.

| Crystal Ligand ID | Trk-A-P1 | Trk-A-P2 | Trk-B-P1 | Trk-B-P2 | Consensus Rank |
| --- | --- | --- | --- | --- | --- |
| MUJ | 584 | 585 | 697 | 695 | 292 |
| LTI | 666 | 431 | 567 | 532 | 589 |
| L0M | 341 | 281 | 249 | **62** | 1193 |
| FQM | 164 | 165 | 120 | **69** | 1161 |
| FQJ | 638 | 691 | 636 | 699 | 574 |
| FQG | **82** | 454 | **2** | **9** | 1277 |
| DZC | **41** | **89** | 125 | **17** | 377 |
| 6UE | **93** | **92** | **29** | **29** | 1245 |
| 6K4 | 273 | 202 | **84** | 244 | 1110 |
| 6K2 | 221 | 262 | **61** | 191 | 1186 |
| 6K1 | 246 | **71** | 244 | 306 | 219 |
| 4EK | 531 | 286 | 340 | 177 | 1142 |
| 4EJ | 326 | **94** | 248 | 235 | 864 |
| 31W | 547 | 539 | 456 | 445 | 995 |
| 31V | **51** | 130 | **94** | 118 | 305 |

In contrast, previously reported Trk agonists ranked poorly when evaluated independently against these pockets (**Table 2**). These findings underscored a major challenge in identifying agonists using conventional docking metrics alone, as inhibitors tended to dominate the highest-ranked results.

**Table 2.** AutoDock Vina individual and consensus docking ranks of published Trk agonists.

| Trk Agonist, CID# | Trk-A-P1 | Trk-A-P2 | Trk-B-P1 | Trk-B-P2 | Consensus Rank |
| --- | --- | --- | --- | --- | --- |
| 903 | 717 | 912 | 983 | 973 | 287 |
| 1880 | 621 | 416 | 419 | 376 | 1050 |
| 2160 | 392 | 723 | 759 | 833 | 193 |
| 3696 | 865 | 968 | 963 | 922 | 483 |
| 3821 | 1002 | 1026 | 1055 | 1030 | 485 |
| 4980 | 1062 | 1075 | 1112 | 1036 | 570 |
| 5881 | 366 | 567 | 477 | 395 | 856 |
| 542158 | 1021 | 1007 | 1018 | 925 | 793 |
| 676310 | 498 | 291 | 411 | 389 | 527 |
| 2054170 | 897 | 1012 | 977 | 997 | 464 |
| **3050408** | **76** | **257** | **447** | **137** | **125** |
| 5281612 | 515 | 415 | 415 | 645 | 329 |
| 44542342 | 370 | 422 | 478 | 476 | 284 |
| 44610701 | 577 | 427 | 517 | 646 | 314 |
| 56951189 | 331 | 166 | 191 | 413 | 273 |

To address this limitation, principal component analysis (PCA) was performed on docking ranks across the four pockets (Trk-A-P1, Trk-A-P2, Trk-B-P1, Trk-B-P2). The PCA results revealed a strong correlation between inhibitor binding ranks across conserved and specificity pockets of Trk-A and Trk-B (**Figure 3A**). This observation suggested that inhibitors tend to bind similarly across multiple pockets and receptors, consistent with their mechanism of blocking receptor activity.

**Figure 3.**
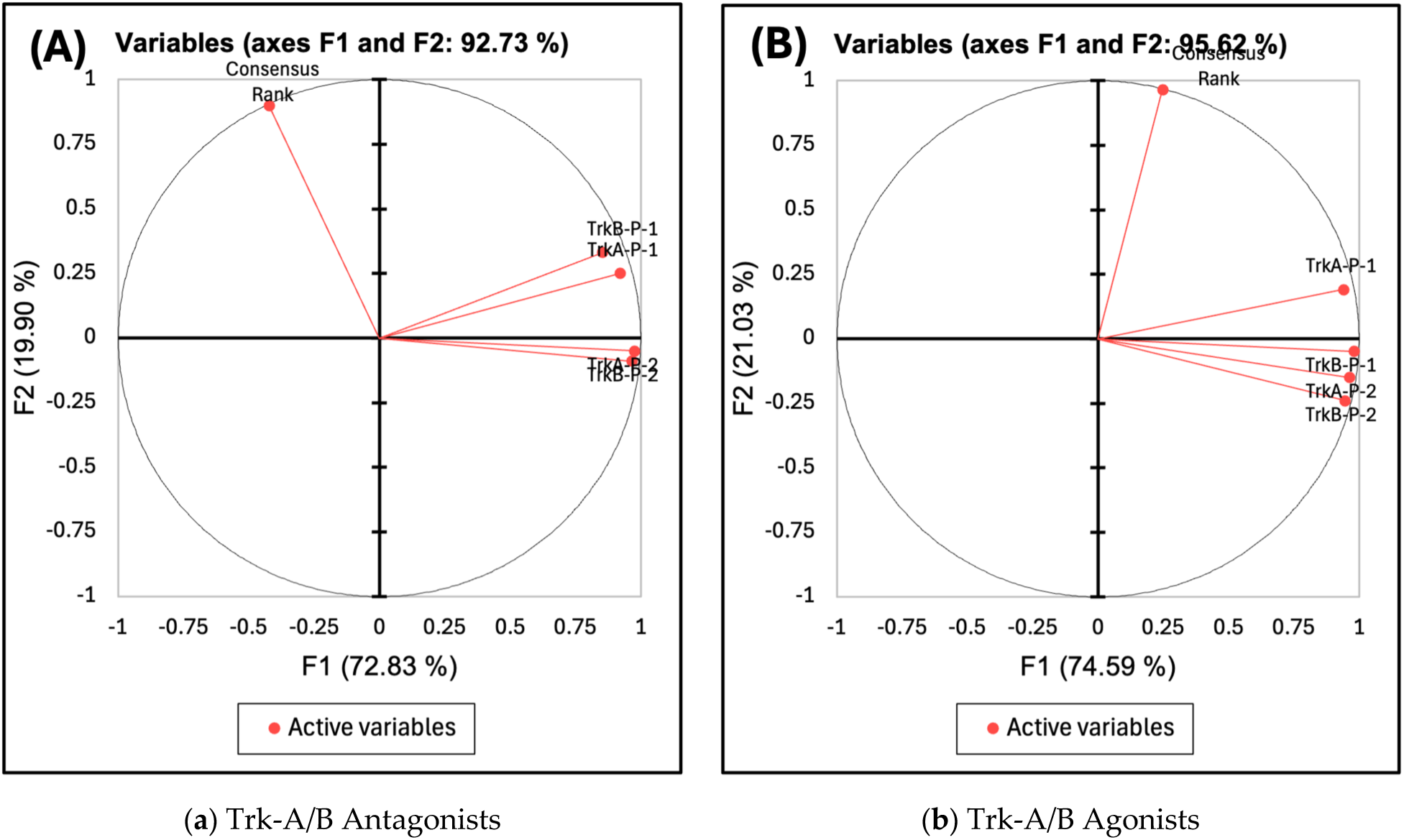
The principal component analysis on the docking data of published (A) antagonists, and (B) agonists of Trk-A and Trk-B receptors.

Based on these insights, we devised a novel consensus-ranking scheme (**Equation 1**) that integrates docking performance across conserved and specificity pockets:

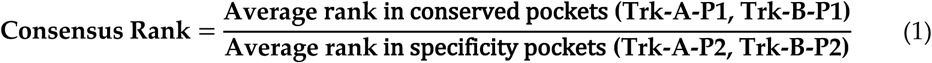

This formulation leveraged the structural similarity of conserved pockets and the diversity of specificity pockets to deprioritize inhibitors while highlighting compounds more likely to act as agonists, with specific emphasis on dual Trk-A and Trk-B agonists. Application of this approach substantially reduced the ranking of known inhibitors, shifting many from top-ranked positions to the bottom 5–10% of the screened dataset (**Table 1**). Most importantly, our novel consensus-ranking scheme improved the positions of several known agonists (**Table 2**). For example, the previously reported dual Trk-A/B agonist Ponazuril (CID_3050408) ranked within the top 10% of hits, whereas other known agonists such as amitriptyline (CID_2160) and LM22B-10 (CID_542158) also showed improved consensus rankings.

Collectively, these analyses demonstrated that our consensus-ranking approach improved discrimination between inhibitors and potential agonists, enabling prioritization of candidate compounds for experimental validation of potential Trk-A/B activation.

### 2.4. Identification of flavonoids as potential dual Trk-A/B agonists

Application of the consensus-ranking approach resulted in a shortlist of 125 compounds comprising approximately the top 10% of screened compounds. Interestingly, most of the top-ranked molecules belonged to steroidal or terpenoid scaffolds, including bufadienolides, alkaloids, and saponins. However, several flavonoid compounds were also identified among the high-ranking hits (**Table 3**).

**Table 3.** The flavonoid compounds from our top hitlist.

| PubChem CID# | Name | Consensus Rank |
| --- | --- | --- |
| 484588 | Euchrestaflavanone A | 18 |
| **14236566** | **Bavachin** | **32** |
| 5316097 | Corylin | 40 |
| 480854 | 3-Hydroxyglabrol | 48 |
| 480865 | (+)-Licoricidin | 53 |
| 5319000 | Licoflavone A | 61 |
| 5320053 | Neobavaisoflavone | 63 |
| 196831 | Licorisoflavan A | 79 |
| 21722008 | Chrysin 6-C-arabinoside 8-C-glucoside | 89 |
| 5488822 | Icariside Ii | 100 |
| 5317478 | Gancaonin A | 103 |
| 24721113 | (-) Catechin hydrate | 104 |
| **193679** | **Isobavachin** | **107** |
| 25056407 | Corylifol A | 110 |
| 155094 | 6-Prenylnaringenin | 111 |
| **513197** | **Isoxanthohumol** | **639** |
| **1880** | **7,8-dihydroxyflavone** | **1050** |

Because flavonoids have previously been reported to modulate neurotrophic signaling [15–17] and also exhibit structural-stabilization activity toward Apolipoprotein E4 [33], the most important genetic risk factor against Alzheimer’s disease, we prioritized flavonoid hits for further analysis and experimental validation.

The functionality of the Trk-A, Trk-B, and Trk-C experimental assays was first confirmed using endogenous neurotrophins as positive controls. In this context, NGF, NT-4/5, and NT-3 produced robust and dose-dependent activation of Trk-A, Trk-B, and Trk-C receptors, respectively (**Figure 4**), validating the reliability of the assay platform, and established baseline receptor activation profiles for comparison with small-molecule candidates.

**Figure 4.**
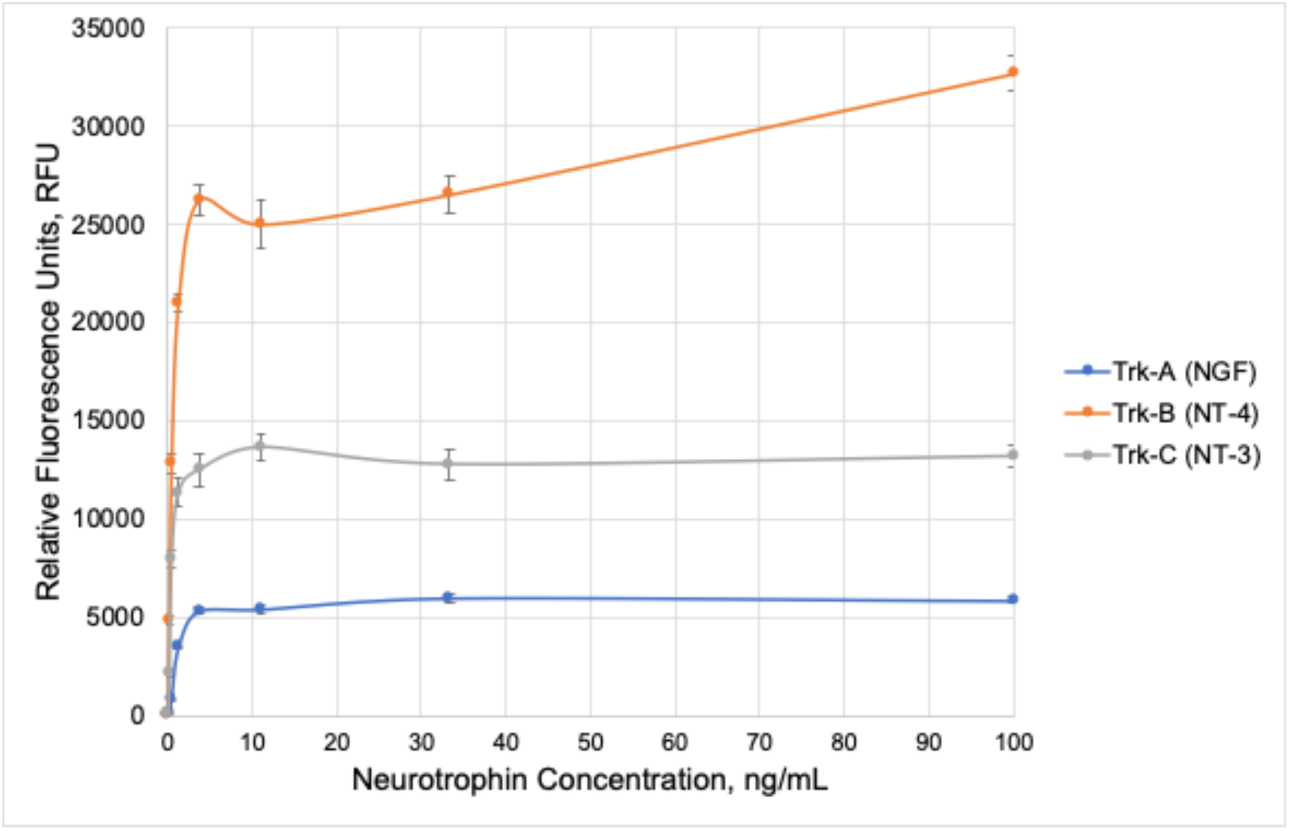
Experimental Trk-A, Trk-B and Trk-C assays: Activation using endogenous ligands (NGF, NT-4/5 and NT-3, respectively).

Among the shortlisted flavonoid compounds, Bavachin (CID_14236566) ranked #32, whereas Isobavachin (CID_193679, consensus rank #107) and Isoxanthohumol (CID_513197, consensus rank #639), the two flavonoids that we had previously discovered as ApoE4 stabilizers [33], were also identified as promising candidates. These compounds, along with the reported Trk-B agonist 7,8-DHF (CID_1880, consensus rank #1050), were selected for their ability to activate Trk receptors. Bavachin demonstrated clear dose-dependent activation of both Trk-A and Trk-B, without Trk-C activation, demonstrating receptor selectivity (**Figure 5**).

**Figure 5.**
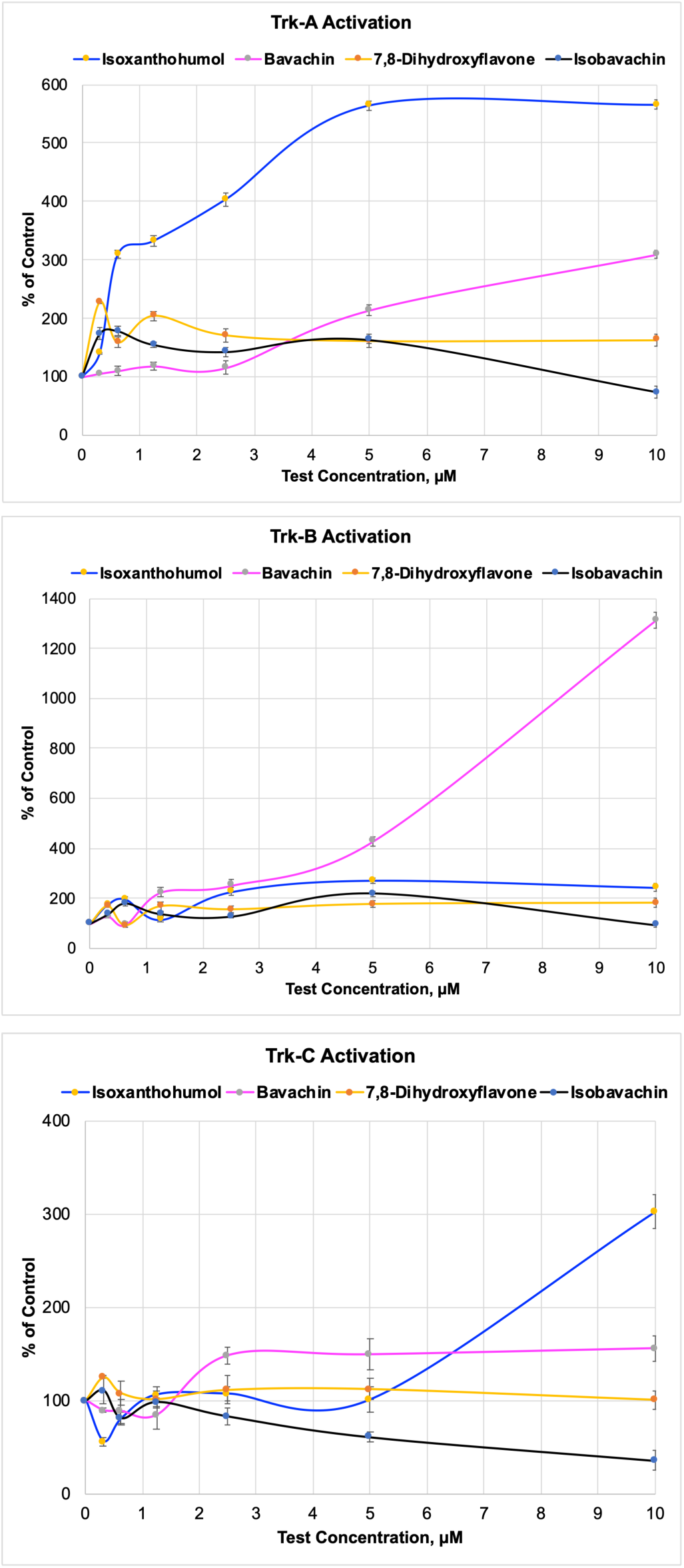
Experimental testing of flavonoid hits in Trk-A, B and C assays.

Moreover, Isoxanthohumol strongly activated Trk-A but showed minimal activity toward Trk-B, and relatively slight activation of Trk-C at only the highest test concentration (10 µM). In contrast, Isobavachin and 7,8-DHF did not activate any of the three receptors under the same experimental conditions.

### 2.5. Binding-mode analysis by induced-fit docking

To understand the molecular basis of receptor activation, induced-fit docking (IFD) was performed for the active flavonoids, as it has been shown to be efficient in handling the receptor flexibility for structurally different ligands [34,35]. In the conserved Trk-A Pocket-1, key interactions observed in the NGF-bound crystal structure, including hydrogen bonding with Asn349 and Gln350 and π-stacking interactions involving Phe327 [22,23], were reproduced by the known agonist Ponazuril (**Figure 6**).

**Figure 6.**
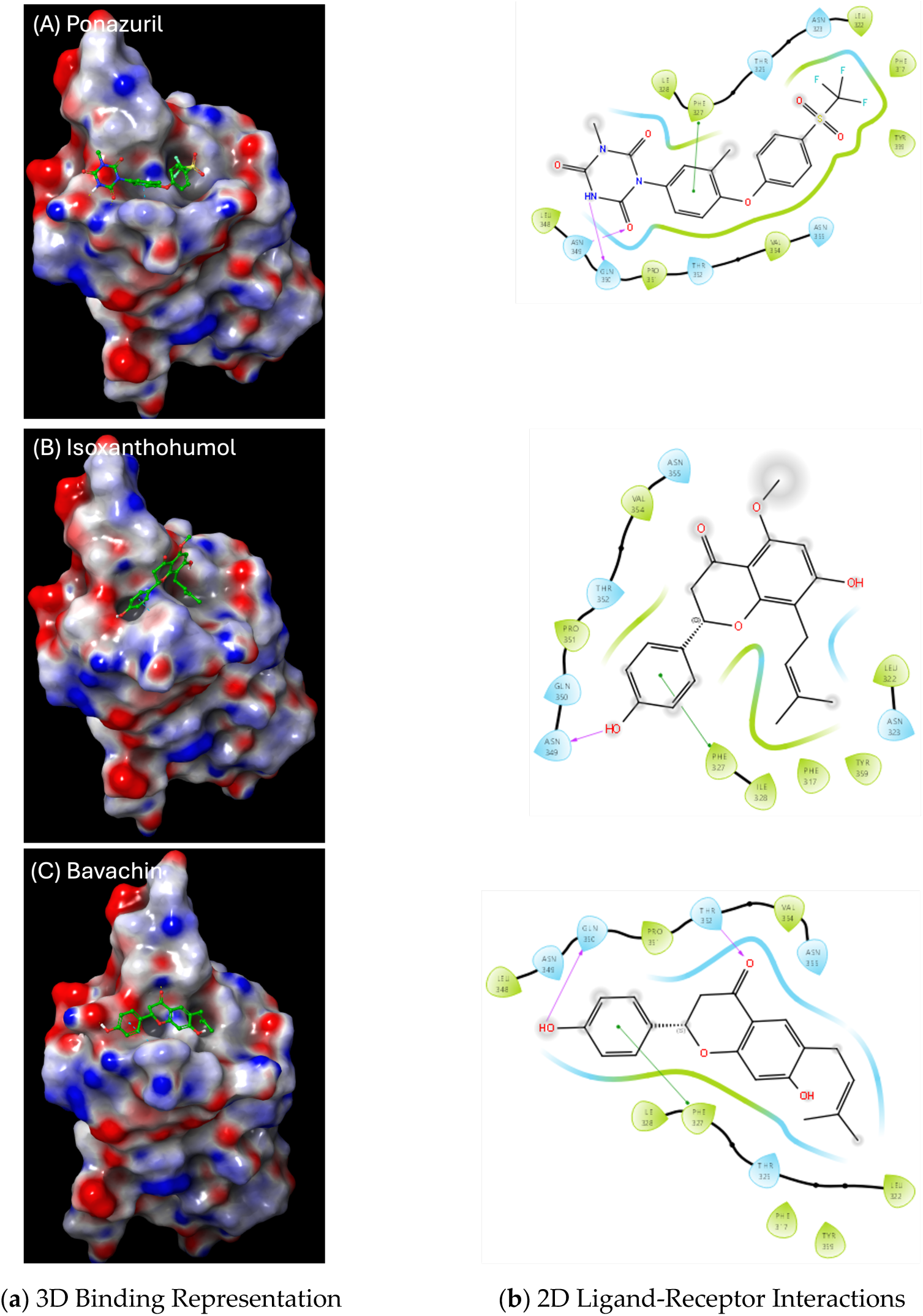
IFD binding modes of top flavonoids against Trk-A-P1: (A) 3D binding representation, (B) 2D ligand-receptor interactions.

Notably, Bavachin closely replicated these interaction features, forming two hydrogen bonds and a π–π interaction with key residues within the pocket, suggesting a binding mode similar to that of Ponazuril. Isoxanthohumol reproduced a subset of these interactions, including hydrogen bonding with Asn349 and aromatic stacking with Phe327, consistent with its experimentally observed Trk-A activation.

In contrast, the specificity pocket of Trk-A is characterized by a hydrophobic canyon formed by residues such as Val294 and Leu333. Isoxanthohumol displayed strong hydrophobic interactions within this pocket (**Figure 7**) and reproduced key hydrogen bonds observed in the NGF-bound Trk-A structure [22], consistent with its strong Trk-A activity observed experimentally.

**Figure 7.**
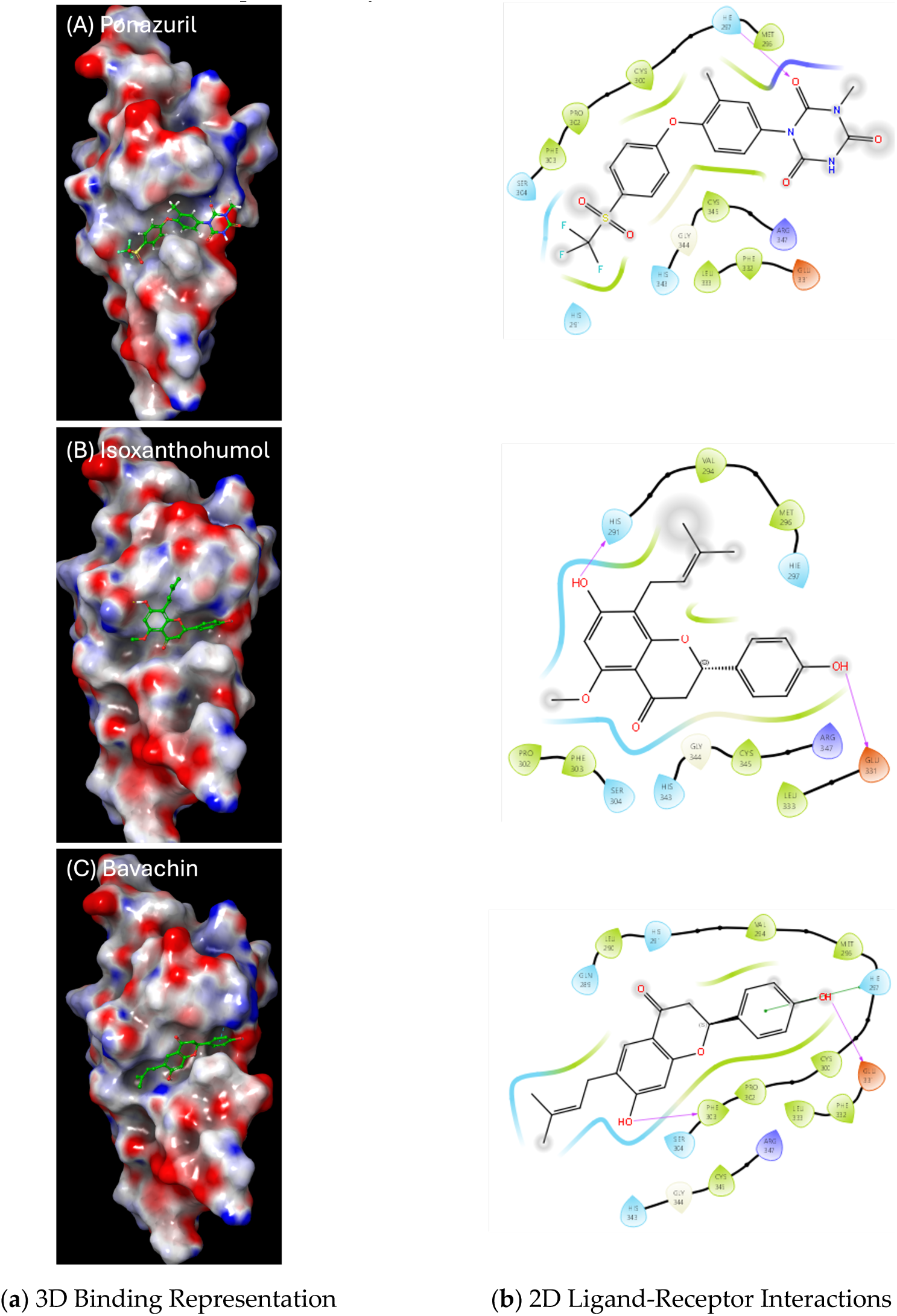
IFD binding modes of top flavonoids against Trk-A-P2: (A) 3D binding representation, (B) 2D ligand-receptor interactions.

For Trk-B, Bavachin formed hydrogen bonds with Asn350 and Thr352 in the conserved pocket (**Figure 8**). The previously reported dual Trk-A/B agonist Ponazuril and Trk-B agonist 7,8-DHF also exhibited H-bond interactions with several key Trk-B amino acids, similar to the endogenous ligand NT-4/5 [24].

**Figure 8.**
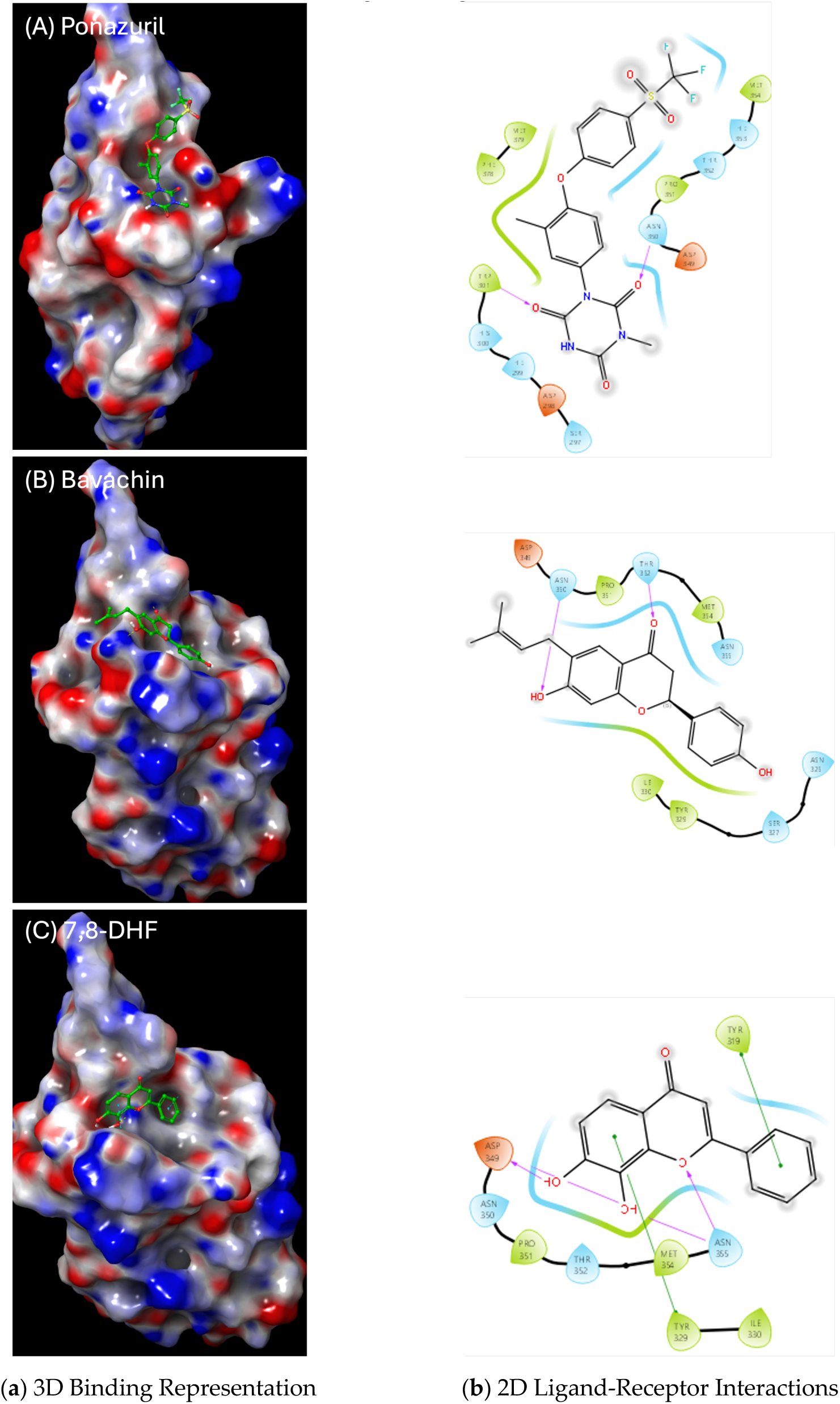
IFD binding modes of top flavonoids against Trk-B-P1: (A) 3D binding representation, (B) 2D ligand-receptor interactions.

In the Trk-B specificity pocket (P-2), NT-4/5 has been shown to form key interactions with several Trk-B receptor amino acids, viz. His-299 (H-bond), Asp-298 (Salt bridge) and His-343 (π–π). The IFD modes of only Ponazuril and 7,8-DHF replicated some of these key interactions, either with these same or nearby residues of Trk-B (**Figure 9**).

**Figure 9.**
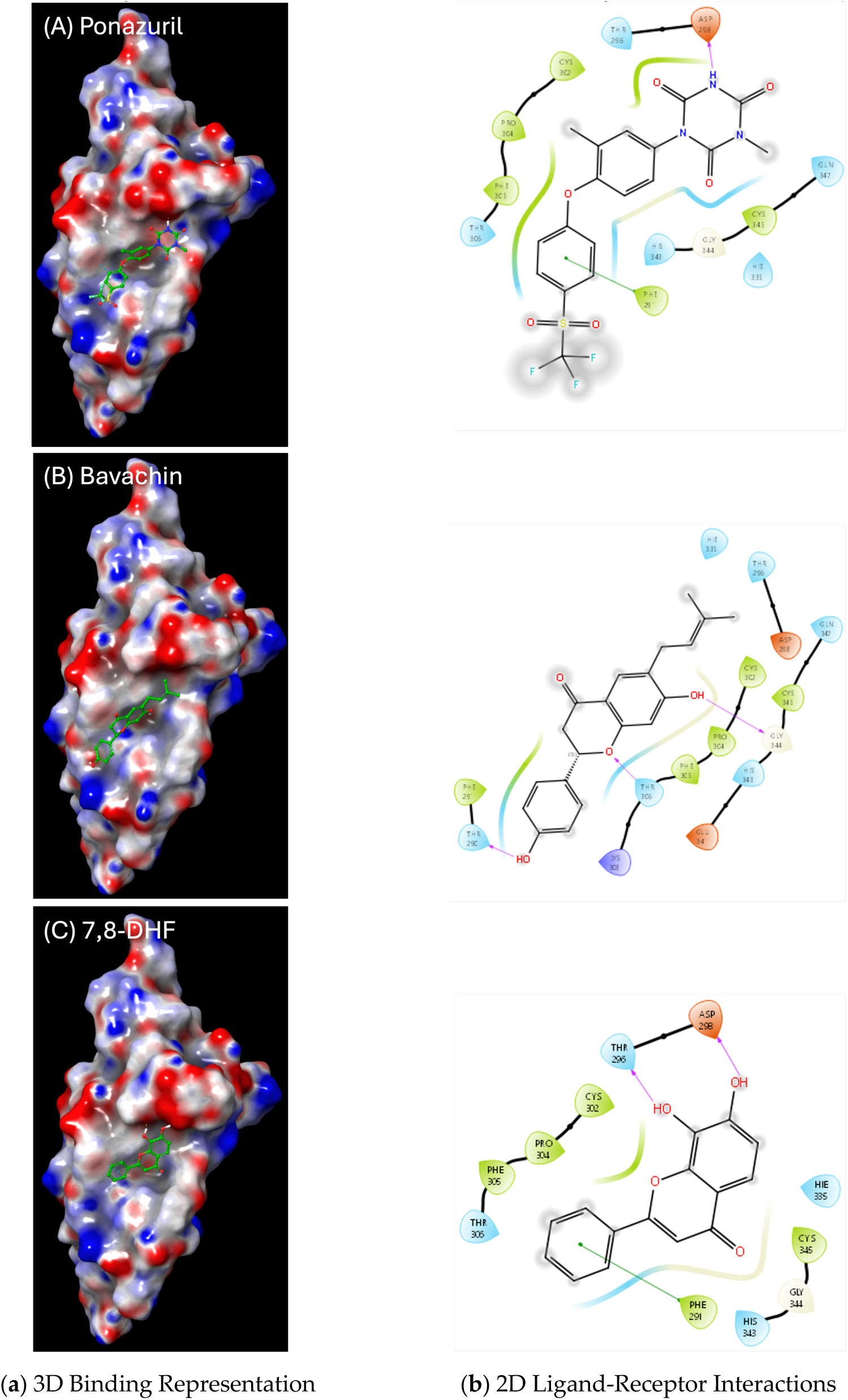
IFD binding modes of top flavonoids against Trk-B-P2: (A) 3D binding representation, (B) 2D ligand-receptor interactions.

### 2.6. Molecular-dynamics simulations reveal Trk-binding stability of active flavonoids

To further validate the docking results, 50-ns molecular-dynamics (MD) simulations were performed for the ligand–receptor complexes to investigate the binding stability of our hits within the Trk-A/B binding pockets. Isoxanthohumol showed highly stable binding in both pockets of Trk-A, consistent with its strong experimental activation of this receptor (**Figure 10A**). In contrast, Isoxanthohumol binding to both Trk-B pockets was relatively highly unstable, as observed by the large ligand RMSD values (data not shown). Bavachin and Ponazuril displayed greater fluctuations within the Trk-A-P1 conserved pocket. In contrast, all three compounds demonstrated stable interactions with the Trk-A-P2 region (**Figure 10B**). The stability of Isoxanthohumol within both Trk-A pockets as opposed to Trk-B may further support its strong Trk-A activation profile observed experimentally.

**Figure 10.**
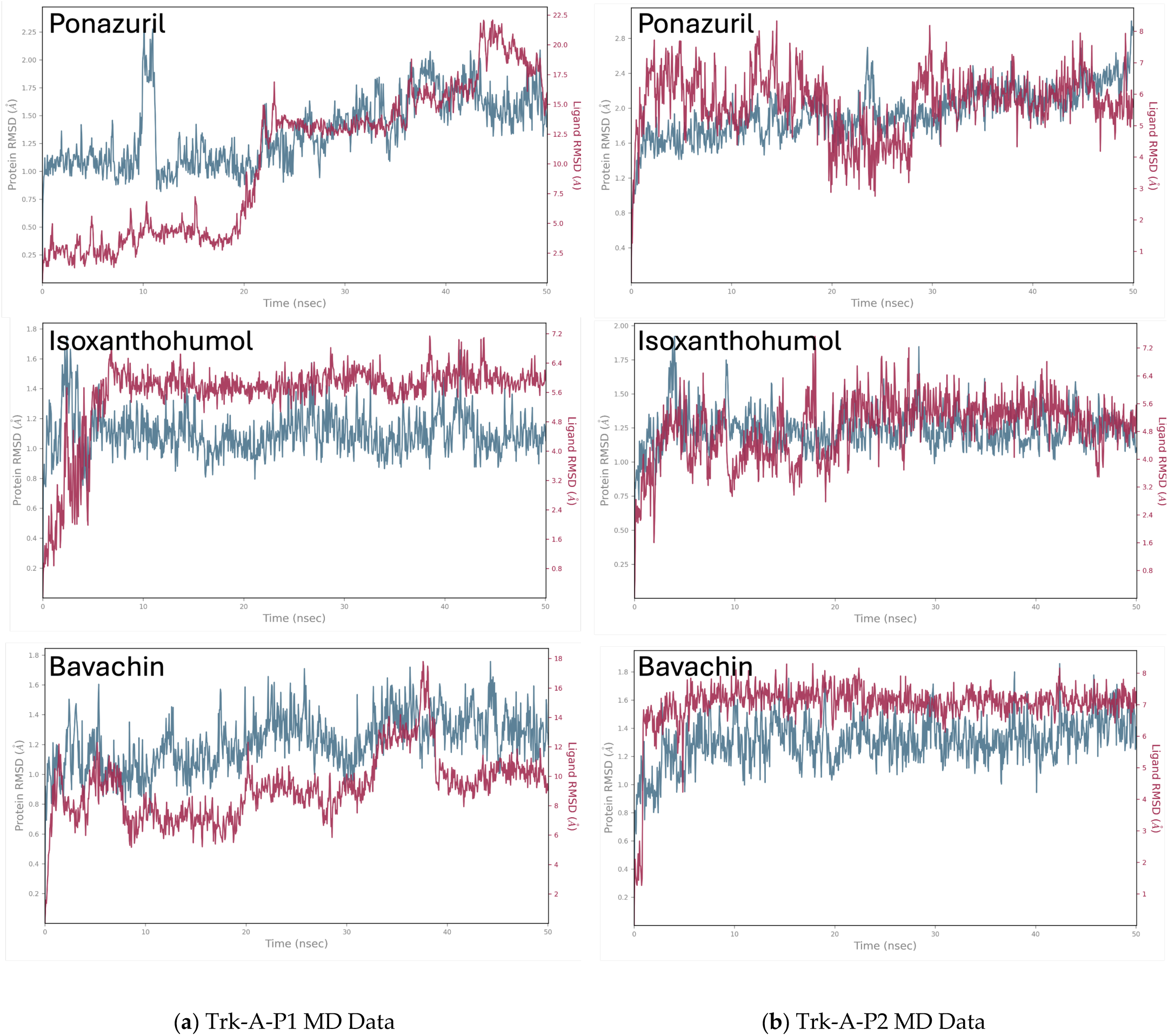
The root-mean-square-distance (RMSD) plots from the MD simulations of flavonoid hits against (A) Trk-A-P1 and (B) Trk-A-P2.

For Trk-B, only Ponazuril maintained relatively stable binding within Trk-B-P1 (**Figure 11A**), while only Bavachin exhibited stable binding within the specificity pocket P-2 (**Figure 11B**). Conversely, 7,8-DHF showed unstable binding across both pockets; these findings are consistent with the lack of Trk-B activation observed experimentally for 7,8-DHF in this study (**Figure 5**).

**Figure 11.**
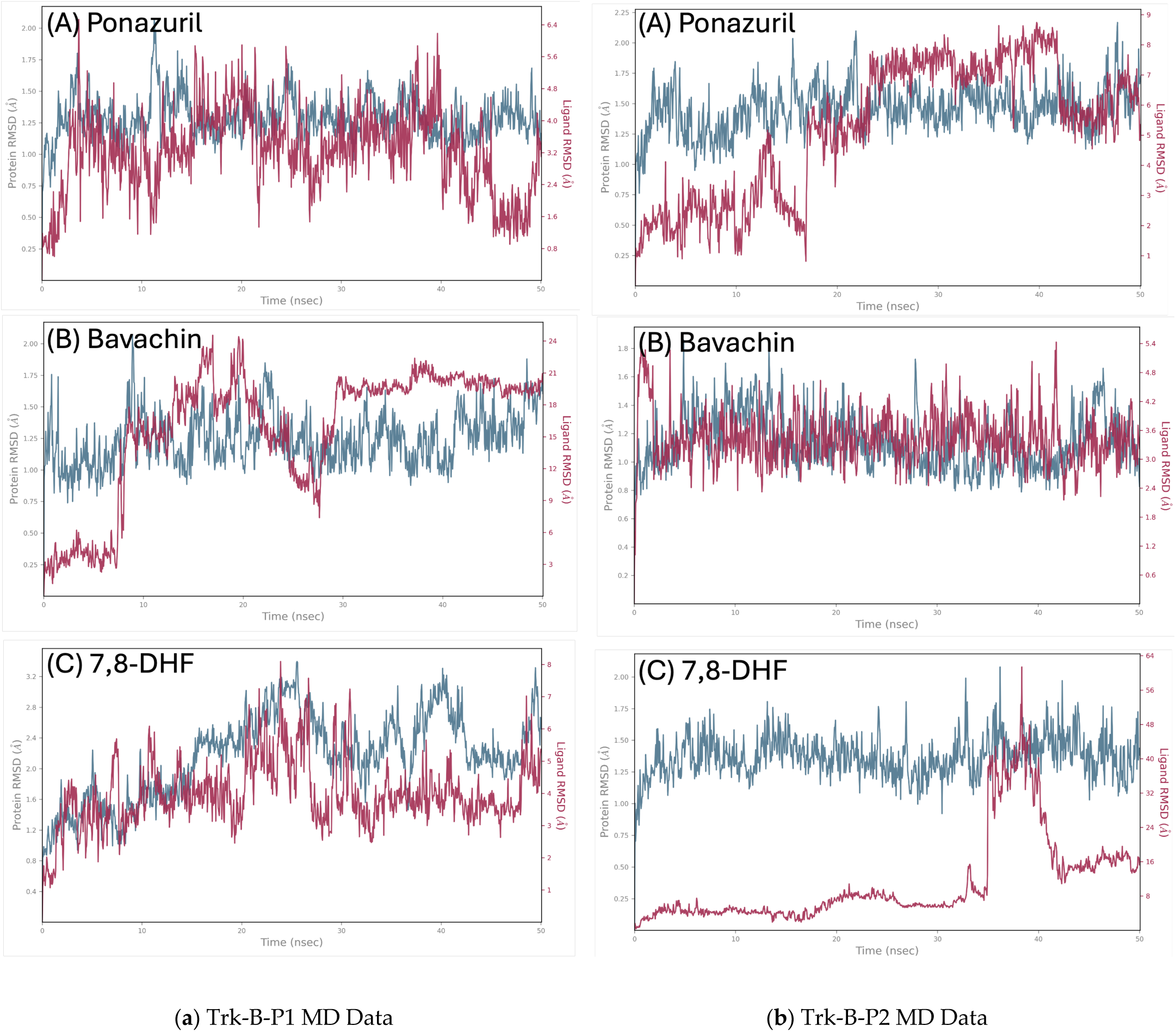
The RMSD plots from the MD simulations of flavonoid hits against (A) Trk-B-P1 and (B) Trk-B-P2.

Taken together, the virtual screening, consensus-ranking analysis, molecular dynamics studies, and experimental validation consistently identified Bavachin as a putative small-molecule neurotrophin mimetic capable of activating both Trk-A and Trk-B receptors. Isoxanthohumol was discovered as a potentially selective Trk-A agonist. In comparison, previously reported Trk-B agonist 7,8-DHF did not show detectable receptor activation against all three Trk receptors, specifically in our current assay conditions.

To further elucidate the structural requirements for Trk-A and Trk-B activation, we performed a systematic SAR comparison of Bavachin with five structurally related analogs (**Figure 12**). Bavachin, characterized by a prenyl group at the C-6 position and a 4’-hydroxyphenyl group, exhibited robust dual agonism. In contrast, Isoxanthohumol, which retains the flavanone core but possesses a methoxy group and a modified prenyl arrangement, demonstrated Trk-A selectivity, failing to activate Trk-B. Interestingly, Isobavachin, a constitutional isomer of Bavachin where the prenyl group is positioned at C-8 instead of C-6, was completely inactive against both receptors. This highlights the critical role of the C-6 prenyl substitution in facilitating the specific hydrophobic interactions required for receptor engagement. Notably, 6-Prenylnaringenin shares a near-identical scaffold with Bavachin; given their structural homology, it is hypothesized to possess similar dual-agonistic potential, though this remains to be experimentally confirmed. Furthermore, the lack of activity observed for 7,8-Dihydroxyflavone in our assays suggests that the flavonoid B-ring hydroxylation pattern and the absence of a bulky hydrophobic prenyl moiety may limit its efficacy in this specific reporter system. Finally, a comparison with the previously reported Isocoumarin analogue (Isocoumarin-1), a known Trk-B agonist in primary hippocampal neurons [36], reveals that while the isocoumarin scaffold can maintain agonism without a prenyl group, the flavanone-based scaffold (as seen in Bavachin) likely relies heavily on the C-6 prenyl tail to compensate for its distinct topological footprint within the Trk-A/B binding pockets.

**Figure 12.**
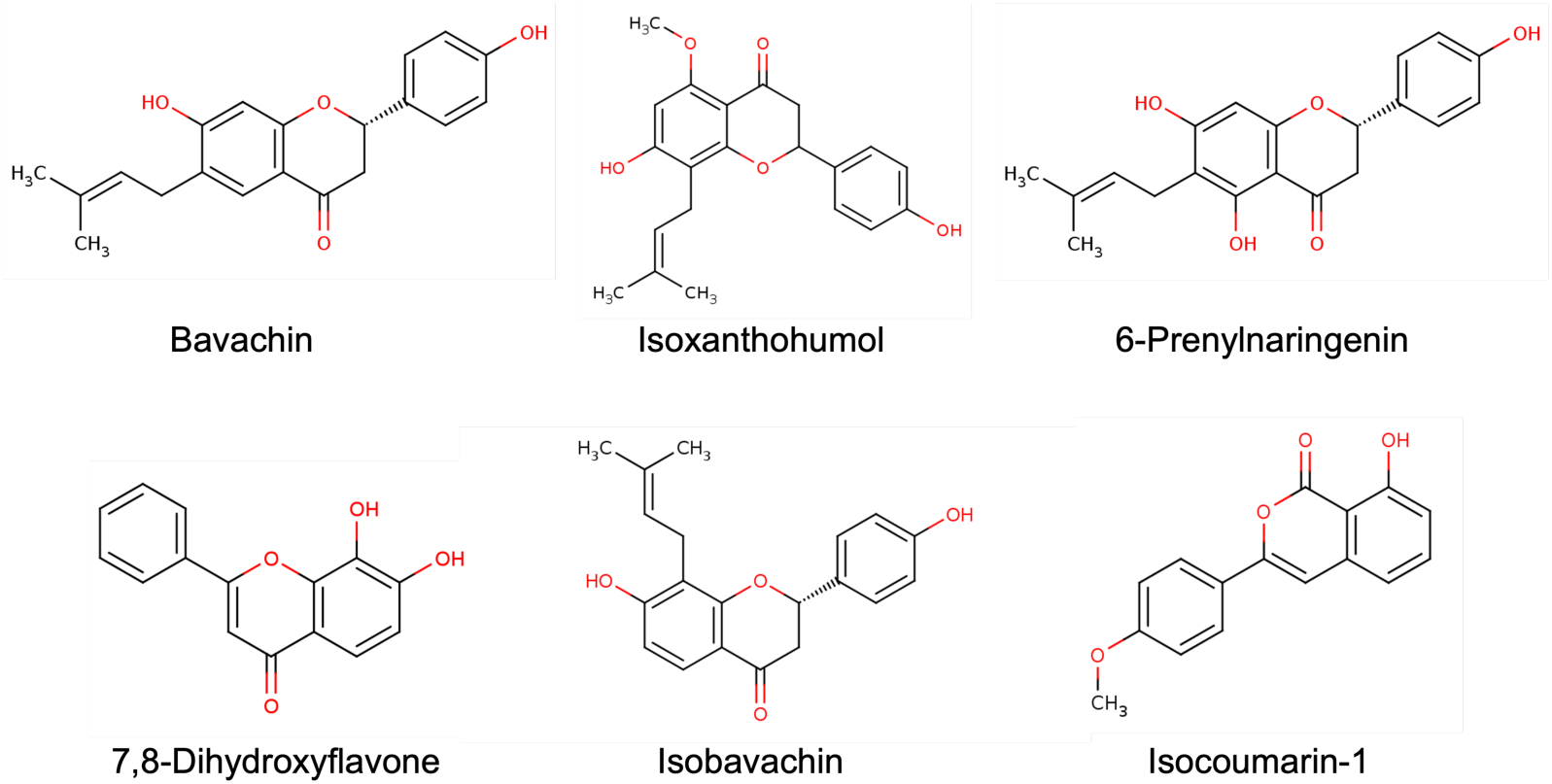
The 2-D chemical structures of Bavachin and other related flavonoids.

## 3. Discussion

The present study integrates structure-based virtual screening, a novel consensus-ranking strategy, and cell-based experimental validation to identify Bavachin as a small-molecule neurotrophin mimetic capable of dual activation of Trk-A and Trk-B receptors. These findings support our central hypothesis that small molecules engaging conserved and receptor-specific neurotrophin-binding pockets can mimic endogenous ligand-induced receptor activation. These findings have significant implications for the development of neuroprotective therapies for Alzheimer’s, Parkinson’s, and other neurodegenerative disorders involving Trk dysregulation. Importantly, this work addresses a long-standing challenge in the field in terms of the reliable identification of true small-molecule Trk agonists through rational drug design efforts as opposed to the more commonly identified inhibitors [14,24].

A key insight from this study is the potential structural basis underlying dual Trk-A/B activation. Previous crystallographic and biochemical studies of Trk–neurotrophin complexes have demonstrated that neurotrophin binding involves coordinated interactions across conserved and receptor-specific variable binding regions of Trk extracellular domains, ultimately promoting receptor dimerization and trans-autophosphorylation [22–24,37]. Our computational analyses indicate that Bavachin may form stabilizing hydrogen-bond and aromatic interactions with key conserved residues involved in endogenous neurotrophin recognition, including Asn349, Gln350, and Phe327 in Trk-A and the corresponding conserved residues Asp349 and Asn350 in Trk-B [22–24]. These findings are consistent with established models of Trk activation, where ligand-induced conformational stabilization drives downstream signaling through PI3K/Akt, MAPK/ERK, and PLCγ pathways [1,2,5,37]. Notably, Bavachin exhibited stable binding within the Trk-B specificity pocket over extended MD simulations, a feature that correlated strongly with its robust experimental activation of Trk-B.

The identification of Bavachin as a potential dual Trk-A and Trk-B agonist is particularly significant in the context of prior efforts to develop small-molecule neurotrophin mimetics. While early studies reported compounds such as 7,8-dihydroxyflavone (7,8-DHF) as selective Trk-B agonists with neuroprotective properties [13,18], subsequent investigations employing multiplexed quantitative assays, including direct receptor binding, dimerization assays, cellular assays, and *in vivo* investigations, have demonstrated that 7,8-DHF fails to induce direct receptor phosphorylation or downstream signaling across various environments [20,21,38]. Our experimental findings, which show a lack of Trk-B activation by 7,8-DHF, are consistent with these later reports and highlight the necessity of combining computational predictions with direct functional assays. This discrepancy also underscores the broader challenge of distinguishing true receptor agonists from compounds that produce downstream phenotypic effects without direct receptor engagement. In this context, MD simulations revealed highly unstable binding of 7,8-DHF in both the conserved (P-1) and specificity (P-2) pockets of Trk-B, further supporting its observed lack of activity in our reporter assays and prior reports questioning its direct Trk-B agonism [20,21,38]. In contrast, Bavachin exhibited stable binding within the Trk-B specificity pocket (P-2) and produced strong receptor activation, suggesting it may be a more reliable scaffold for dual Trk-A and Trk-B agonist development. Our computational data provide a potential structural rationale suggesting that 7,8-DHF may lack the topological complementarity required for stable orthosteric engagement of the Trk-B D5 domain, further supporting the emerging consensus that its neuroprotective effects may be mediated through Trk-independent mechanisms.

A key methodological contribution of this work is the development of a consensus-ranking approach that integrates docking performance across conserved and specificity pockets of both Trk-A and Trk-B receptors. Principal component analysis revealed that known Trk inhibitors tend to bind similarly across these pockets, whereas agonists display more heterogeneous and receptor-specific interaction profiles. By integrating docking performance across complementary binding sites, our novel consensus-ranking scheme effectively deprioritized inhibitor-like compounds while enriching for agonist-like candidates. Application of this strategy successfully elevated known Trk agonists, including Ponazuril and Amitriptyline, and led to the identification of Bavachin among top-ranked computational hits. These results highlight the importance of multi-site, receptor-aware scoring frameworks for agonist discovery, particularly for extracellular receptor tyrosine kinases whose activation depends on ligand-induced conformational rearrangements rather than simple active-site occupancy [14,25,39,40]. It is noteworthy here that we have previously devised and utilized a similar consensus-ranking scheme involving apo- and holo-pockets on Apolipoprotein E4 (ApoE4), the most significant genetic risk factor for Alzheimer’s disease [33]. That approach led to the identification of Isobavachin as a potential structure corrector and stabilizer of ApoE4, demonstrating the utility of this multi-pocket ranking system in identifying bioactive flavonoids for neurodegenerative targets.

From a pharmacological standpoint, the enrichment of flavonoids among the top hits is consistent with a growing body of literature highlighting their neuroprotective and cognition-enhancing properties, mediated through antioxidant effects, modulation of intracellular signaling, and regulation of synaptic plasticity [15–17,41]. In addition, several flavonoids have demonstrated the ability to cross the blood–brain barrier and exert central nervous system effects, making them attractive scaffolds for neurotherapeutic development [41,42]. Bavachin, in particular, may offer added advantages due to its natural origin and potential bioavailability, as natural products derived from traditional medicine systems are often enriched for favorable pharmacokinetic and safety profiles [26,27].

The differential activity observed among structurally related flavonoids further provides insight into structure–activity relationships governing Trk activation. Isoxanthohumol exhibited selective Trk-A activation, consistent with its stable binding within Trk-A pockets and instability in Trk-B, while Isobavachin showed no detectable activity against all Trk receptors. It is noteworthy here that both Isobavachin and Isoxanthohumol, but not Bavachin, were recently discovered by us as structure correctors of Apolipoprotein E4 (ApoE4) [33], the most significant genetic risk factor of Alzheimer’s disease. These findings suggest that subtle variations in functional groups and binding orientation can significantly influence receptor specificity and efficacy. Similar structure-dependent modulation of neurotrophic signaling has been reported for other flavonoid derivatives, reinforcing the importance of precise molecular interactions in determining biological activity [41].

In the broader context of neurodegenerative disease therapeutics, the dual activation of Trk-A and Trk-B receptors may provide synergistic benefits. Impaired NGF/Trk-A signaling contributes to degeneration of basal forebrain cholinergic neurons, while reduced BDNF/Trk-B signaling is associated with synaptic dysfunction and cognitive decline in Alzheimer’s disease and other neurodegenerative disorders [6–9,43]. Therapeutic strategies that simultaneously target multiple neurotrophic pathways are increasingly recognized as more effective than single-target approaches for complex diseases such as Alzheimer’s disease [44–47]. In this regard, our newly discovered putative dual Trk-A/B agonist Bavachin represents a promising lead compound with unique therapeutic advantages, which may help modulate multiple aspects of neuronal survival and function.

While the present study provides compelling evidence for the interaction between Bavachin and the Trk-A/B receptors, we acknowledge that our experimental validation relies primarily on engineered cell-based reporter assays. These systems are highly sensitive and provide an excellent platform for high-throughput screening and initial validation of receptor activation; however, they do not fully replicate the physiological complexity of endogenous neuronal environments.

Consequently, in light of the current experimental scope, we characterize Bavachin as a putative dual Trk-A/Trk-B agonist. We recognize that the leap from reporter activity to definitive neurotrophin mimicry requires further biochemical confirmation. Future studies are necessary to provide direct evidence of receptor engagement through Western blot analysis or ELISA-based quantification of specific Trk phosphorylation sites (e.g., p-TrkA Tyr490 and p-TrkB Tyr516/Tyr706/707). Furthermore, evaluating the activation of classic downstream signaling cascades, such as the PI3K/AKT, MAPK/ERK and CREB phosphorylation pathways, in native or neuronal-like systems (e.g., PC12 or SH-SY5Y cells) will be essential to confirm that Bavachin-induced signaling translates into functional biological responses.

Furthermore, we recognize that a key hallmark of functional neurotrophin mimetics is their ability to translate the neurotrophin receptor activation into complex phenotypic outcomes, such as neurite outgrowth, axonal branching, and neuronal survival. Although phenotypic assays in models such as PC12 cells or primary hippocampal neurons were beyond the scope of this initial discovery-phase report, they remain the ‘gold standard’ for confirming biological mimicry. It is possible that small-molecule agonists may trigger receptor phosphorylation or reporter activity without achieving the specific spatio-temporal signaling required for full morphological differentiation—a phenomenon sometimes observed in non-functional Trk transactivation. Consequently, Bavachin should currently be considered a potent lead Trk-A/B agonist. Future studies should focus on evaluating the neuroprotective effects of Bavachin in native neuronal systems to fully validate its therapeutic potential as a functional neurotrophin mimetic.

A significant question remains regarding how a monomeric flavonoid like Bavachin induces the requisite dimerization of Trk receptors, a process typically driven by large dimeric proteins such as NGF or BDNF. We propose that Bavachin may function as a ‘molecular glue’ agonist. Molecular glues are small molecules that bind at protein-protein interfaces, lowering the entropic barrier for dimerization and stabilizing the active complex, driving drive proximity-dependent signaling [48-50]. Our computational analysis suggests that Bavachin occupies a hydrophobic pocket at the D5 domain interface, potentially acting as a bridge that stabilizes the Trk-A or Trk-B homodimer in an active state. However, as our current evidence relies on *in silico* stability and reporter-based functional signaling, further biophysical studies using Co-Immunoprecipitation (Co-IP) or Bioluminescence Resonance Energy Transfer (BRET) are warranted to definitively characterize the stoichiometry and kinetics of Bavachin-induced potential receptor dimerization. Elucidating such precise mechanism of receptor activation, whether through direct receptor dimer stabilization, or allosteric modulation, will be critical for guiding rational drug design efforts against this clinically relevant target.

Finally, the clinical translation of flavonoid-based leads is frequently hampered by rapid first-pass metabolism and restricted systemic bioavailability. Previous investigations into the pharmaco-kinetic (PK) profile of Bavachin have noted that it is subject to extensive metabolic conversion, resulting in peak plasma concentrations below 10 ng/mL [55]. However, despite these low systemic levels, tissue distribution studies have confirmed that Bavachin effectively penetrates the blood-brain barrier, reaching therapeutic targets within cerebral nuclei [56]. This suggests that the Bavachin possesses a high intrinsic propensity for CNS entry, likely facilitated by its optimal lipophilicity. To overcome the challenges of metabolic instability, future development of Bavachin as a neurotherapeutic may require advanced drug delivery strategies, such as the synthesis of prodrug derivatives or the implementation of nanoparticle-mediated delivery systems. Such optimizations will be essential to transition Bavachin from a potent *in vitro* lead to a viable *in vivo* neuroprotective agent.

In summary, our present data combined with the proposed studies may further help establish the therapeutic potential of Bavachin in the context of neurodegenerative pathology.

## 4. Materials and Methods

### Structure-Based Virtual Screening

The DCABM-TCM database was prepared for docking by extracting SMILES from: http://bionet.ncpsb.org.cn/dcabm-tcm/ [28]. The database included ∼1,740 bioavailable TCM compounds detected in the blood, out of which ∼ 500 molecules with high molecular weights were removed. We included 15 each agonists and antagonists of the Trk receptors that are previously published, in our screening database (DCABM-TCM). The 3D conformers for this combined database were generated using OMEGA 6.1.1.1 3D conformer generator [51] from OpenEye, Cadence Molecular Sciences, Santa Fe, NM, http://www.eyesopen.com. The ensemble docking of this drug database was carried out against two each pockets on the Trk-A (PDB: 1WWW) and Trk-B (PDB: 1HCF) proteins downloaded from the protein data bank (PDB) [52]. Binding pockets were identified based on ligand-bound crystal structures and further analyzed using the DoGSiteScorer tool to evaluate druggability scores and geometric features. These Trk protein structures were then processed using the AutoDock Tools utility [53], whereby bound neurotrophin ligands and water molecules were removed, polar hydrogens were added, non-polar hydrogens were merged, and Gasteiger charges were assigned for all the atoms in the proteins. The Trk proteins and ligand atoms were converted into PDBQT format. The AutoDock Vina docking algorithm [32] was then used to carry out structure-based docking of the DCABM-TCM molecules to the conserved (P-1) and specificity (P-2) pockets on Trk-A and Trk-B proteins. The Trk-A and Trk-B structures were overlapped and thus the same search space coordinates were used for docking into their two pockets: P-1 [Center- X: 9.2105, Y: -2.5016, Z: 4.5091; Size- X: 25 Å, Y: 25 Å, Z: 25 Å], and P-2 [Center- X: 19.7446, Y: -2.3479, Z: -12.9946; Size- X: 25 Å, Y: 25 Å, Z: 25 Å]. Default docking parameters were used, and the docked ligands were ranked according to their best docking score values. Principal Component Analysis (PCA) was performed on docking rank datasets to identify correlations between binding across multiple pockets. A novel consensus-ranking scheme utilizing docking ranks against both the Trk-A and Trk-B pockets was used to shortlist top ∼10% of the molecules (125 hits) for further analysis.

### Consensus-Ranking Strategy

To identify high-confidence candidates for dual Trk-A/Trk-B activation, a consensus-ranking scheme was designed to integrate structural and energetic data across multiple binding sites while mitigating the scoring bias inherent in any single pocket. Prior to screening, a preliminary druggability analysis was conducted using the DoGSiteScorer server, which identified high-volume, ‘druggable’ pockets overlapping with the endogenous neurotrophin-binding interfaces. These regions were categorized as either ‘conserved’ (sharing high homology between Trk-A and Trk-B) or ‘specificity’ pockets (exhibiting significant divergence in amino acid composition and physicochemical characteristics).

Our selection hypothesis was grounded in the premise that a true dual agonist must maintain high topological complementarity across the conserved domains of both receptors. Conversely, we posited that compounds scoring highly only within ‘specificity’ pockets were more likely to represent false positives or selective binders rather than the intended dual-acting mimetics. To operationalize this, we devised a consensus-ranking formula (Equation 1):

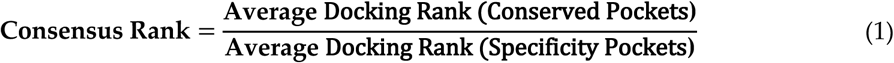

In this framework, a low numerator (indicating high affinity for conserved sites) combined with a higher denominator (indicating lower relative affinity for specificity sites) yields a lower overall consensus rank. This mathematical arrangement effectively enriches the hit list for potential dual-agonistic scaffolds, while deprioritizing compounds with profiles more characteristic of site-specific inhibitors.

### Induced-Fit Docking (IFD)

Selected compounds were subjected to induced-fit docking to account for receptor flexibility and optimize ligand–receptor interactions. Specifically, the flexible-receptor docking of top structure-based virtual screening hits was carried out within both the conserved and specificity pockets of Trk-A (PDB ID: 1WWW) and Trk-B (PDB ID: 1HCF) receptors. The IFD protocol was run with default parameters from the Schrodinger’s Maestro graphical user interface. The first stage of IFD protocol involved extended sampling using Glide docking algorithm to generate 80 initial poses through softened-potential docking step, which allows tolerance of more steric clashes as compared to a normal, rigid-receptor docking protocol. The second stage then involved protein refinement using Prime module to allow the active site conformational change around 5 Å of the initial ligand poses. Herein, the energy minimization was carried out using OPLS4 force field and Prime’s implicit solvent model. Finally, the active site was further refined by Prime followed by redocking by Glide XP scoring function leading to the generation of the final IFD models. The top 5 IFD poses for each molecule were graphically visualized using Maestro and the best conformation for each molecule based on the observed binding mode and the IFD score was chosen for further analysis including molecular dynamics (MD) simulations as described below.

### Molecular Dynamics Studies

The binding stability of our top hits was investigated using the molecular dynamics (MD) simulations of the respective ligand-protein complexes using Schrodinger’s Desmond MD module as described previously [54]. Briefly, the top binding conformations from the induced-fit docking were chosen as the starting conformations for the MD simulations. The ligand interactions were modeled with the OPLS4 force field. The ligand-protein docking complexes were solvated using TIP4P-EW water model in an orthorhombic boundary box of 10 Å size in all three directions, followed by their neutralization with appropriate number of Cl-ions. Also, the salt was added at 0.15 M concentration to simulate the physiological salt concentration. Default parameters in Desmond including OPLS4 force field were utilized to equilibrate this molecular system. Finally, each of the equilibrated systems was subjected to 50 ns of MD simulations using the NPT ensemble class to maintain a constant temperature of 300 K and a constant pressure of 1 atm. Temperature control was achieved using the Nosé-Hoover thermostat, while the Martyna-Tobias-Klein barostat was employed to regulate pressure. The atom coordinates were recorded at every 50 ps for the follow-up simulation analyses. The root mean square deviation (RMSD) for the proteins and the docked ligands were calculated over the entire simulation trajectory with reference to their respective first frames. The MD protocol employed a sequential sampling strategy (3 x 50 ns). Specifically, the stability of top hits was verified using a multi-stage approach where the final coordinate frame of each run served as the starting configuration for the subsequent simulation. This method was chosen to enhance phase-space sampling and ensure the reproducibility of the observed binding modes, a strategy our group has previously validated across targets such as ApoE4, P53-MDM2 and the PD-1/PD-L1 axis [33,35,39].

### Reagents and Compounds

Bavachin and other flavonoid analogs were purchased from [Cayman Chemicals (Ann Arbor, MI, USA) and MedChemExpress, (Monmouth Junction, NJ, USA)]. The chemical purity of all tested compounds was verified to be ≥98% as determined by Liquid Chromatography-Mass Spectrometry (LC-MS) and provided by the manufacturer’s certificate of analysis.

### Trk Receptor Reporter Assays

These assays utilize engineered mammalian cells expressing the respective Trk receptor and a luciferase reporter, where receptor activation triggers intracellular signaling leading to luminescence proportional to receptor activity. Assay-ready cryopreserved cells were thawed and plated in 96-well plates using the supplied assay medium according to the manufacturer’s instructions. The chemical purity of all tested compounds was verified to be ≥98% as determined by High-Performance Liquid Chromatography (HPLC) and provided by the manufacturer’s certificate of analysis. Test compounds were prepared in dimethyl sulfoxide (DMSO) and diluted to the desired concentration, with a final DMSO concentration of ≤0.1% (v/v). Cells were treated in a dose-dependent concentration manner in triplicate. Vehicle controls (DMSO) and positive controls, viz. nerve growth factor (NGF) for Trk-A, NT-4/5 for Trk-B and NT-3 for Trk-C, were included for assay validation and normalization. Following incubation at 37 °C with 5% CO2 for 24 h, luciferase detection reagent was added, and luminescence was measured using BioTek Synergy Neo microplate reader. Data were background-corrected and normalized to vehicle controls to determine fold activation.

### Statistical Analysis

Results are reported as mean ± standard deviation from three independent experiments. Statistical significance was determined using one-way Analysis of Variance (ANOVA) followed by Tukey’s post-hoc test for multiple comparisons. P-values < 0.05 were considered statistically significant.

## 5. Conclusions

In summary, our present study identified Bavachin as a putative dual Trk-A and Trk-B agonist and established a generalizable integrated computational–experimental framework for discovering small-molecule neurotrophin mimetics. Future studies focused on downstream signaling, *in vivo* efficacy, and structure–activity optimization of the Bavachin scaffold may provide a strong foundation for the development of novel, orally bioavailable therapeutics for the treatment of neurodegenerative disorders.

## Institutional Review Board Statement

Not applicable.

## Informed Consent Statement

Not applicable.

## Data Availability Statement

The data are available on request from the corresponding author.

## Acknowledgments

The authors thank John Stoddart (Computer Science, Widener University) for his kind help with the GPU-enabled molecular dynamics simulations. The authors acknowledge Olivia Allred’s and Eugenia Khanales’ role as undergraduate researchers and Ishaan Joshi’s role as a high school student intern learning molecular docking in our laboratory.

## References

1. Huang, E.J.; Reichardt, L.F. Neurotrophins: Roles in neuronal development and function. Annu. Rev. Neurosci. 2001, 24, 677–736. 10.1146/annurev.neuro.24.1.677

2. Kaplan, D.R.; Miller, F.D. Neurotrophin signal transduction in the nervous system. Curr. Opin. Neurobiol. 2000, 10, 381–391. 10.1016/S0959-4388(00)00092-1

3. Chao, M.V. Neurotrophins and their receptors: A convergence point for many signalling pathways. Nat. Rev. Neurosci. 2003, 4, 299–309. 10.1038/nrn1078

4. Barbacid, M. The Trk family of neurotrophin receptors. J. Neurobiol. 1994, 25, 1386–1403. 10.1002/neu.480251107

5. Segal, R.A. Selectivity in neurotrophin signaling: Theme and variations. Annu. Rev. Neurosci. 2003, 26, 299–330. 10.1146/annurev.neuro.26.041002.131421

6. Allen, S.J.; Watson, J.J.; Dawbarn, D. The neurotrophins and their role in Alzheimer’s disease. Curr. Neuropharmacol. 2011, 9, 559–573. 10.2174/157015911798376190

7. Zuccato, C.; Cattaneo, E. Brain-derived neurotrophic factor in neurodegenerative diseases. Nat. Rev. Neurol. 2009, 5, 311–322. 10.1038/nrneurol.2009.54

8. Nagahara, A.H.; Tuszynski, M.H. Potential therapeutic uses of BDNF in neurological and psychiatric disorders. Nat. Rev. Drug Discov. 2011, 10, 209–219. 10.1038/nrd3366

9. Counts, S.E.; Mufson, E.J. The role of nerve growth factor receptors in cholinergic basal forebrain degeneration in prodromal Alzheimer disease. J. Neuropathol. Exp. Neurol. 2005, 64, 263–272. 10.1093/jnen/64.4.263

10. Thoenen, H.; Sendtner, M. Neurotrophins: From enthusiastic expectations through sobering experiences to rational therapeutic approaches. Nat. Neurosci. 2002, 5, 1046–1050. 10.1038/nn938

11. Poduslo, J.F.; Curran, G.L. Permeability at the blood–brain and blood–nerve barriers of the neurotrophic factors: NGF, CNTF, NT-3, BDNF. Brain Res. Mol. Brain Res. 1996, 36, 280–286. 10.1016/0169-328x(95)00250-v

12. Massa, S.M.; Yang, T.; Xie, Y.; Shi, J.; Bilgen, M.; Joyce, J.N.; Nehama, D.; Rajadas, J.; Longo, F.M. Small molecule BDNF mimetics activate TrkB signaling and prevent neuronal degeneration in rodents. J. Clin. Invest. 2010, 120, 1774–1785. 10.1172/JCI41356

13. Jang, S.W.; Liu, X.; Yepes, M.; Shepherd, K.R.; Miller, G.W.; Liu, Y.; Wilson, W.D.; Xiao, G.; Blanchi, B.; Sun, Y.E.; et al. A selective TrkB agonist with potent neurotrophic activities by 7,8-dihydroxyflavone. Proc. Natl. Acad. Sci. USA 2010, 107, 2687–2692. 10.1073/pnas.0913572107

14. Longo, F.M.; Massa, S.M. Small-molecule modulation of neurotrophin receptors: A strategy for the treatment of neurological disease. Nat. Rev. Drug Discov. 2013, 12, 507–525. 10.1038/nrd4024

15. Spencer, J.P. Flavonoids and brain health: multiple effects underpinned by common mechanisms. Genes Nutr. 2009, 4, 243–250. 10.1007/s12263-009-0136-3

16. Williams, R.J.; Spencer, J.P.E. Flavonoids, cognition, and dementia: Actions, mechanisms, and potential therapeutic utility. Free Radic. Biol. Med. 2012, 52, 35–45. 10.1016/j.freeradbiomed.2011.09.010

17. Vauzour, D. Dietary polyphenols as modulators of brain functions: Biological actions and molecular mechanisms underpinning their beneficial effects. Oxid. Med. Cell Longev. 2012, 2012, 914273. 10.1155/2012/914273

18. Liu, X.; Chan, C.B.; Jang, S.W.; Pradoldej, S.; Huang, J.; He, K.; Phun, L.H.; France, S.; Xiao, G.; Jia, Y.;, et al. A synthetic 7,8-dihydroxyflavone derivative promotes neurogenesis and exhibits potent antidepressant effect. J. Med. Chem. 2010, 53, 8274–8286. 10.1021/jm101206p

19. Andero, R.; Heldt, S.A.; Ye, K.; Liu, X.; Armario, A.; Ressler, K.J. Effect of 7,8-dihydroxyflavone, a small-molecule TrkB agonist, on emotional learning. Am. J. Psychiatry 2011, 168, 163–172. 10.1176/appi.ajp.2010.10030326

20. Boltaev, U.; Meyer, Y.; Tolibzoda, F.; Jacques, T.; Gassaway, M.; Xu, Q.; Wagner, F.; Zhang, Y.L.; Palmer, M.; Holson, E.; Sames, D. Multiplex quantitative assays indicate a need for reevaluating reported small-molecule TrkB agonists. Sci. Signal. 2017, 10, eaal1670. 10.1126/scisignal.aal1670

21. Todd, D.; Gowers, I.; Dowler, S.J.; Wall, M.D.; McAllister, G.; Fischer, D.F.; Dijkstra, S.; Fratantoni, S.A.; van de Bospoort, R.; Veenman-Koepke, J.; Flynn, G.; Arjomand, J.; Dominguez, C.; Munoz-Sanjuan, I.; Wityak, J.; Bard, J.A. A monoclonal antibody TrkB receptor agonist as a potential therapeutic for Huntington’s disease. PLoS ONE 2014, 9, e87923. 10.1371/journal.pone.0087923

22. Wiesmann, C.; Ultsch, M.H.; Bass, S.H.; de Vos, A.M. Crystal structure of nerve growth factor in complex with the ligand-binding domain of the TrkA receptor. Nature 1999, 401, 184–188. 10.1038/43705

23. Banfield, M.J.; Naylor, R.L.; Robertson, A.G.; Allen, S.J.; Dawbarn, D.; Brady, R.L. Specificity in Trk receptor: Neurotrophin interactions: The crystal structure of TrkB-d5 in complex with neurotrophin-4/5. Structure 2001, 9, 1191–1199. 10.1016/s0969-2126(01)00681-5

24. Ferreira, L.G.; dos Santos, R.N.; Oliva, G.; Andricopulo, A.D. Molecular docking and structure-based drug design strategies. Molecules 2015, 20, 13384–13421. 10.3390/molecules200713384

25. Ultsch, M.H.; Wiesmann, C.; Simmons, L.C.; Henrich, J.; Yang, M.; Reilly, D.; Bass, S.H.; de Vos, A.M. Crystal structures of the neurotrophin-binding domain of TrkA, TrkB and TrkC. J. Mol. Biol. 1999, 290, 149–159. 10.1006/jmbi.1999.2816

26. Newman, D.J.; Cragg, G.M. Natural products as sources of new drugs from 1981 to 2014. J. Nat. Prod. 2016, 79, 629–661. 10.1021/acs.jnatprod.5b01055

27. Huang, L.; Xie, D.; Yu, Y.; Liu, H.; Shi, Y.; Shi, T.; Wen, C. TCMID 2.0: A comprehensive resource for traditional Chinese medicine. Nucleic Acids Res. 2018, 46, D1117–D1120. 10.1093/nar/gkx1028

28. Liu, X.; Liu, J.; Fu, B.; Chen, R.; Jiang, J.; Chen, H.; Li, R.; Xing, L.; Yuan, L.; Chen, X.;, et al. DCABM-TCM: A database of constituents absorbed into the blood and metabolites of traditional Chinese medicine. J. Chem. Inf. Model. 2023, 63, 4948–4959. 10.1021/acs.jcim.3c00365

29. Patil, S.P.; Maki, S.; Khedkar, S.A.; Rigby, A.C.; Chan, C. Withanolide A and asiatic acid modulate multiple targets associated with amyloid-beta precursor protein processing and amyloid-beta protein clearance. J. Nat. Prod. 2010, 73, 1196–1202. 10.1021/np900633j

30. Patil, S.P.; Tran, N.; Geekiyanage, H.; Liu, L.; Chan, C. Curcumin-induced upregulation of the anti-tau cochaperone BAG2 in primary rat cortical neurons. Neurosci. Lett. 2013, 554, 121–125. 10.1016/j.neulet.2013.09.008

31. Volkamer, A.; Kuhn, D.; Rippmann, F.; Rarey, M. DoGSiteScorer: A web server for automatic binding site prediction, analysis and druggability assessment. Bioinformatics 2012, 28, 2074–2075. 10.1093/bioinformatics/bts310

32. Trott, O.; Olson, A.J. AutoDock Vina: Improving the speed and accuracy of docking with a new scoring function, efficient optimization, and multithreading. J. Comput. Chem. 2010, 31, 455–461. 10.1002/jcc.21334

33. Patil, S.P.; Kuehn, B.R.; McCullough, C.; Bates, D.; Hazim, H.; Diallo, M.; Francois, N. Discovery of isobavachin, a natural flavonoid, as an apolipoprotein E4 (ApoE4) structure corrector for Alzheimer’s disease. Molecules 2025, 30, 940. 10.3390/molecules30040940

34. Sherman, W.; Day, T.; Jacobson, M.P.; Friesner, R.A.; Farid, R. Novel procedure for modeling ligand/receptor induced fit effects. J. Med. Chem. 2006, 49, 534–553. 10.1021/jm050540c

35. Patil, S.P.; Pacitti, M.F.; Gilroy, K.S.; Ruggiero, J.C.; Griffin, J.D.; Butera, J.J.; Notarfrancesco, J.M.; Tran, S.; Stoddart, J.W. Identification of antipsychotic drug fluspirilene as a potential p53-MDM2 inhibitor: A combined computational and experimental study. J. Comput. Aided Mol. Des. 2015, 29, 155–163. 10.1007/s10822-014-9811-6

36. Sudarshan, K.; Boda, A.K.; Dogra, S.; Bose, I.; Yadav, P.N.; Aidhen, I.S. Discovery of an isocoumarin analogue that modulates neuronal functions via neurotrophin receptor TrkB. Bioorg. Med. Chem. Lett. 2019, 29, 585–590. 10.1016/j.bmcl.2018.12.057

37. Lemmon, M.A.; Schlessinger, J. Cell signaling by receptor tyrosine kinases. Cell 2010, 141, 1117–1134. 10.1016/j.cell.2010.06.011

38. Pankiewicz, P.; Szybiński, M.; Kisielewska, K.; Gołębiowski, F.; Krzemiński, P.; Rutkowska-Włodarczyk, I.; Moszczyński-Pętkowski, R.; Gurba-Bryśkiewicz, L.; Delis, M.; Mulewski, K.;, et al. Do small molecules activate the TrkB receptor in the same manner as BDNF? Limitations of published TrkB low molecular agonists and screening for novel TrkB orthosteric agonists. Pharmaceuticals 2021, 14, 704. 10.3390/ph14080704

39. Patil, S.P.; Ballester, P.J.; Kerezsi, C.R. Prospective virtual screening for novel p53-MDM2 inhibitors using ultrafast shape recognition. J. Comput. Aided Mol. Des. 2014, 28, 89–97. 10.1007/s10822-014-9732-4

40. Fattakhova, E.; Hofer, J.; DiFlumeri, J.; Cobb, M.; Dando, T.; Romisher, Z.; Wellington, J.; Oravic, M.; Radnoff, M.; Patil, S.P. Identification of the FDA-approved drug pyrvinium as a small-molecule inhibitor of the PD-1/PD-L1 interaction. ChemMedChem 2021, 16, 2769–2774. 10.1002/cmdc.202100264

41. Spencer, J.P. The impact of flavonoids on memory: physiological and molecular considerations. Chem. Soc. Rev. 2009, 38, 1152–1161. 10.1039/b800422f

42. Youdim, K.A.; Qaiser, M.Z.; Begley, D.J.; Rice-Evans, C.A.; Abbott, N.J. Flavonoid permeability across an in situ model of the blood-brain barrier. Free Radic. Biol. Med. 2004, 36, 592–604. 10.1016/j.freeradbiomed.2003.11.023

43. Budni, J.; Bellettini-Santos, T.; Mina, F.; Garcez, M.L.; Zugno, A.I. The involvement of BDNF, NGF and GDNF in aging and Alzheimer’s disease. Aging Dis. 2015, 6, 331–341. 10.14336/ad.2015.0825

44. Gong, C.-X.; Dai, C.-L.; Liu, F.; Iqbal, K. Multi-targets: An unconventional drug development strategy for Alzheimer’s disease. Front. Aging Neurosci. 2022, 14, 837649. 10.3389/fnagi.2022.837649

45. Pasieka, A.; Panek, D.; Malawska, B. Multifunctional ligand approach: Search for effective therapy against Alzheimer’s disease. In Alzheimer’s Disease: Drug Discovery; Exon Publications: Brisbane, Australia, 2020; pp. 1–25.

46. Crunkhorn, S. Combination therapy alleviates AD-related pathologies. Nat. Rev. Drug Discov. 2025, 24, 742. 10.1038/d41573-025-00146-3

47. Cheong, S.L.; Tiew, J.K.; Fong, Y.H.; Leong, H.W.; Chan, Y.M.; Chan, Z.L.; Kong, E.W.J. Current pharmacotherapy and multi-target approaches for Alzheimer’s disease. Pharmaceuticals 2022, 15, 1560. 10.3390/ph15121560

48. Fischer, E.S.; Böhm, K.; Lydeard, J.R.; Yang, H.; Stadler, M.B.; Cavadini, S.; Nagel, J.; Serluca, F.; Acker, V.; Lingaraju, G.M.; Tichkule, R.B.; Schebesta, M.; Forrester, W.C.; Schirle, M.; Hassiepen, U.; Ottl, J.; Hild, M.; Beckwith, R.E.; Harper, J.W.; Jenkins, J.L.; Thomä, N.H. Structure of the DDB1-CRBN E3 ubiquitin ligase in complex with thalidomide. Nature 2014, 512, 49–53. 10.1038/nature13527

49. Schreiber, S.L. The rise of molecular glues. Cell 2021, 184, 3–9. 10.1016/j.cell.2020.12.020

50. Schlessinger, J. Receptor tyrosine kinases: Legacy of the first two decades. Cold Spring Harb. Perspect. Biol. 2014, 6, a008912. 10.1101/cshperspect.a008912

51. Hawkins, P.C.D.; Skillman, A.G.; Warren, G.L.; Ellingson, B.A.; Stahl, M.T. Conformer generation with OMEGA: Algorithm and validation using high-quality structures from the Protein Data Bank and the Cambridge Structural Database. J. Chem. Inf. Model. 2010, 50, 572–584. 10.1021/ci100031x

52. Berman, H.M.; Battistuz, T.; Bhat, T.N.; Bluhm, W.F.; Bourne, P.E.; Burkhardt, K.; Feng, Z.; Gilliland, G.L.; Iype, L.; Jain, S.;, et al. The Protein Data Bank. Acta Crystallogr. D Biol. Crystallogr. 2002, 58, 899–907. 10.1107/S0907444902003451

53. Morris, G.M.; Huey, R.; Lindstrom, W.; Sanner, M.F.; Belew, R.K.; Goodsell, D.S.; Olson, A.J. AutoDock4 and AutoDockTools4: Automated docking with selective receptor flexibility. J. Comput. Chem. 2009, 30, 2785–2791. 10.1002/jcc.21256

54. DiFrancesco, M.; Hofer, J.; Aradhya, A.; Rufinus, J.; Stoddart, J.; Finocchiaro, S.; Mani, J.; Tevis, S.; Visconti, M.; Walawender, G.;, et al. Discovery of small-molecule PD-1/PD-L1 antagonists through combined virtual screening and experimental validation. Comput. Biol. Chem. 2023, 102, 107804. 10.1016/j.compbiolchem.2022.107804

55. Gao, Q.; Xu, Z.; Zhao, G.; Wang, H.; Weng, Z.; Pei, K.; Wu, L.; Cai, B.; Chen, Z.; Li, W. Simultaneous quantification of five main components of Psoralea corylifolia L. in rat plasma by ultra-high-performance liquid chromatography–tandem mass spectrometry. J. Chromatogr. B 2016, 1011, 128–135. 10.1016/j.jchromb.2015.12.044

56. Wang, Y.F.; Zhang, Y.B.; Chen, Z.J.; Zhang, Y.T.; Yang, X.W. Plasma pharmacokinetics and cerebral nuclei distribution of major constituents of Psoraleae fructus in rats after oral administration. Phytomedicine 2018, 38, 166–174. 10.1016/j.phymed.2017.12.002

